# Decoding Spatial Programs in Human Glioblastoma Through Surprisal Information-Theoretical Analysis

**DOI:** 10.64898/2026.06.23.733938

**Authors:** Shuozhen Bao, Gretchen Long, Shouryo Ghose, Nuoya Wang, Haikuo Li, Mei Zhong, Logan Matthews, Yao Lu, Jie Sheng, Zewei Tu, William Escobar, Pallavi Gopal, Declan McGuone, Zeynep E. Erson Omay, Khaled Alok, Di Zhang, Marcello DiStasio, Mircea Tesileanu, Mingyu Yang, Keyi Li, Jennifer Moliterno, James R. Heath, Micha Sam Brickman Raredon, Francoise Remacle, Raphael D. Levine, Jiangbing Zhou, Rong Fan

## Abstract

Glioblastoma (GBM) is a highly heterogeneous and invasive brain tumor in which complex interactions among tumor cells, immune cells, and neurons shape disease progression and therapeutic resistance. Resolving spatial patterns programs in glioblastoma (GBM) requires analytical approaches that go beyond variance-driven embeddings. Here, we applied thermodynamic surprisal analysis, an information-theoretic decomposition framework, to spatial transcriptomic sequencing from human GBM specimens to identify orthogonal constraint modes that capture dominant and previously hidden spatial programs. Surprisal analysis revealed structured patterns in the data that are not highlighted by standard approaches such as PCA or cell type deconvolution, including immune-activation–associated signatures as well as spatial programs consistent with synapse remodeling. Coupling surprisal decomposition with spatial ligand–receptor interaction analysis, NICHES, along with multiplexed protein imaging connected these spatial hidden modes to reveal potential communication networks. Together, these results position surprisal analysis as a powerful, complementary strategy for interrogating spatial tumor architecture, enabling discovery of non-obvious spatial programs and interactions that are obscured by variance-based methods.

## Introduction

Glioblastoma (GBM), the most malignant (CNS grade 4) astrocytoma, remains a uniformly lethal disease despite aggressive surgical resection, radiation, and temozolomide therapy, with a 5-year survival rate of only 6.9% and a recurrence rate approaching 90%^1, 2^. As the most common malignant brain tumor in adults, accounting for 54% of all gliomas and 16% of all primary brain tumors, GBM displays extraordinary inter- and intra-tumoral heterogeneity that underlies its treatment resistance and dismal clinical outcomes^3^. Although single-cell sequencing has revealed diverse cellular states and remarkable plasticity^4–8^, it remains inadequately understood how GBM evolves, adapts, and reorganizes itself across the complex spatial landscape of the human brain.

Glioblastoma (GBM) is increasingly understood not merely as a genetically heterogeneous malignancy but as a disease organized around distinct spatially defined microenvironmental niches that shape tumor identity, plasticity, and clinical behavior^9–11^. Spatial omics studies now reveal that GBM comprises several recurring ecological structures including neuron–tumor, vascular/perivascular, invasive edge, and hypoxic or necrotic microenvironments, each exerting specific regulatory pressures on tumor cells^12–14^. High-resolution spatial profiling has mapped endothelial–tumor and pericyte–tumor interactions, extracellular matrix remodeling, hypoxia-induced angiogenic programs, and gradients of stemness and therapy resistance along perivascular spaces^12, 14–17^. These niches also exhibit strong immune exclusion, altered myeloid cell states, and metabolic specialization, supporting longstanding models that position the perivascular niche as a central regulator of glioma stem cell maintenance, metabolic adaptation, and vascular-guided invasion. Nerve–tumor niches have recently emerged as a defining feature of GBM biology^18–25^. Pioneering work in cancer neuroscience has shown that glioma cells form functional synapses with neurons, engage in activity-dependent signaling, and remodel local neural circuit processes that promote tumor proliferation, excitability, and network integration. Spatial transcriptomic and proteomic analyses further identify ligand–receptor signaling, synaptic scaffolding components, and electrophysiologically coupled tumor populations within these niches, underscoring their role as active growth-promoting microenvironments. At the tumor boundaries, invasive edge niches drive GBM’s hallmark infiltrative behavior^9, 26^. Spatial omics and multiomic integration highlight transcriptional programs associated with motility, axonal tract migration, extracellular matrix reorganization, and oxidative or lipid metabolic shifts that facilitate diffuse infiltration. These niches often contain hybrid lineage states with elevated plasticity, representing key contributors to treatment escape and recurrence. Hypoxic and necrotic niches^27^, often aligned with mesenchymal transition states, exhibit oxygen deprivation, inflammatory signaling, myeloid recruitment, metabolic rewiring, and dense extracellular matrix deposition. These niches serve as hubs of immune suppression and therapeutic resistance^28–30^, reinforcing their role as a critical mesenchymal microenvironment within GBM. Together, these neuron–tumor, vascular, invasive, and hypoxic/necrotic niches constitute the core spatial architecture of GBM.

Yet each individual GBM specimen provides only a partial and biased snapshot of this ecosystem: the abundance, arrangement, and dominance of niches vary dramatically across patients, making it difficult to discern which spatial features are foundational and which are patient-specific. As a result, even with advanced spatial transcriptomics and multiplexed imaging, current analyses, typically applied to single tumors in isolation, cannot readily reconstruct the overarching spatial programs of GBM or identify the latent constraints that generate recurrent architectures across tumors. These limitations underscore the need for an analytical framework capable of assembling multiple partial spatial omics observations into a unified cross-tumor representation, revealing shared spatial programs, niches, cell-cell interaction, and the governing principles underlying GBM tumor dynamics, plasticity, and progression.

In this study, we analyzed spatial omics with thermodynamic surprisal analysis^31^, an information-theoretic method^32^ originally developed in statistical physics, as a principled approach for identifying hidden constraint modes that govern GBM spatial organization. Surprisal analysis has previously been applied to GBM^33^ to identify the state of minimal Gibbs free energy^34^ when disease constraints are not imposed.

Using integrated spatial transcriptomic (DBiT-seq) and multiplexed protein imaging across ten spatial regions from 10 paired tumor samples from five IDH-wild-type GBM patients, we reframed each sample as a partial realization of common underlying spatial programs. Surprisal analysis decomposes these data into orthogonal constraint modes, representing deviations from a theoretical steady state, that collectively reconstruct the major spatial processes active across GBM tissues.

By linking surprisal-derived constraint modes with spatial ligand–receptor interactype mapping, we highlight the common signaling networks that define neural niches across multiple tumors—including synaptic L-R interactions such as NLGN1–NRXN1 and SEMA5A–MET that show hijacked neuron behaviors. Together, this work demonstrates that spatial omics surprisal analysis provides an integrated theoretical framework for assembling heterogeneous cell states into a cohesive, generalizable blueprint of human GBM architecture. Rather than analyzing each tumor in isolation, this approach reveals the spatial principles that consistently underlie GBM organization, highlighting small-GTPase activation and neuron–tumor crosstalk as central, recurrent features of disease progression and potential therapeutic vulnerability.

## Results

### Sample Collection, Experimental Workflow, and Surprisal Analysis Framework

We acquired ten GBM specimens from five patients, each comprising a paired T1-contrast–enhanced (Core) and non–contrast-enhanced (Peripheral) region. Samples were immediately snap frozen in OCT embedding using isopentane and liquid nitrogen and stored at −80°C. Cryo-sectioning was performed to obtain 10 μm-thick sections on coated glass slides, with adjacent slides documented. All samples were profiled by Patho-DBiT spatial transcriptomics (n = 10) and six samples were additionally profiled by PhenoCycler multiplexed protein imaging (n = 6), providing 25 μm spatial resolution across 2.5 mm × 2.5 mm tissue areas (Fig. 1A). The ten regions from five patients spanned patients ages 48 to 74 years, collected from both male and female patients. All samples are IDH wild-type with TERT promoter mutations (Supplemental Table 1, Supplemental Figure 1).

**Figure 1.**
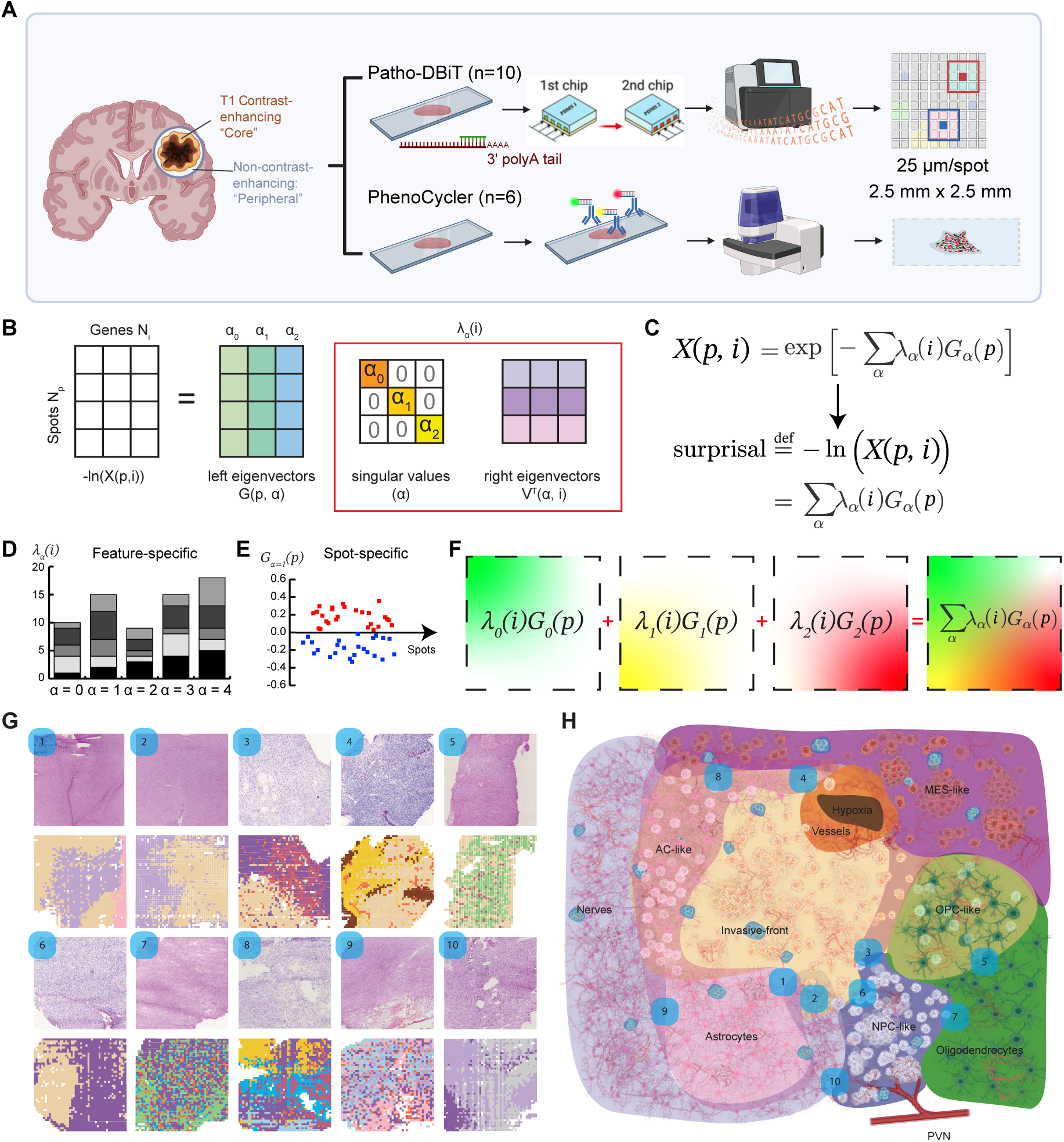
Spatial multi-omics profiling and spatial information-theoretic surprisal analysis framework for GBM. (A) Experimental workflow. Ten GBM specimens from five patients, each comprising a paired T1-contrast–enhanced (Core) and non–contrast-enhanced (Peripheral) region, were profiled by Patho-DBiT spatial transcriptomics (n = 10) and PhenoCycler multiplexed protein imaging (n = 6), with 25 μm spatial resolution across 2.5 mm × 2.5 mm tissue areas. (B) Schematic of the surprisal analysis decomposition. The natural-log-transformed expression matrix − ln[*X*(*p*, *i*)] is factored by singular value decomposition (SVD) into left eigenvectors *G_α_*(*p*)(spatial amplitudes), singular values, *α* (constraint magnitudes), and right eigenvectors, *λ_α_*(*i*) (gene contributions), where *p* indexes spatial spots and *i* indexes genes. (C) Definition of the gene-by-spot expression matrix *X*(*p*, *i*). (D) Illustration of feature-specific gene contributions, *λ_α_*(*i*) and spot-specific spatial amplitudes *G_α_*(*p*). Singular value magnitudes decrease with increasing constraint index α, reflecting the diminishing contribution of higher-order modes. (E-F) Cumulative reconstruction of spatial gene expression. Progressive summation of constraint terms *G*_0_(*p*)*λ*_0_(*i*)+ *G*_1_(*p*)*λ*_1_(*i*) + *G*_2_(*p*)*λ*_2_(*i*) + … converges toward the observed expression profile, with spatial maps of individual constraint amplitudes (*G*_1_, *G*_2_, *G*_3_) shown. (G) Sample information and phenotype characterization of all samples. (H) Schematic of GBM heterogeneity and tumor microenvironment based on our spatial samples.

Tools like Seurat^35^ and Scanpy^36^ use Principal Component Analysis for initial data reduction, a statistical technique for simplifying data complexity by transforming original variables into a smaller set of principal components which capture the most variance. This allows reduction of high-dimensional gene expression data, making it easier to identify and visualize distinct cell populations. Conversely, surprisal analysis measures the unexpectedness of events based on information theory and entropy, providing a quantitative measure of how surprising an observation is against expected probabilities. While PCA focuses on variance and dimensionality reduction, surprisal analysis focuses on the thermodynamic deviation from steady state and informational content of data. Surprisal analysis^31^ is particularly useful in heterogeneous and complex contexts, like GBM^33^, and we chose this method to provide deeper insights into the unpredictability and dynamics of cellular behavior in more challenging datasets such as those affected by disease-driven transcriptional plasticity.

Surprisal analysis was developed to analyze how systems in non-equilibrium deviate from a steady state due to the action of constraints, internal and environmental. It was applied to a wide range of chemical^32^, physical^37, 38^ and biological systems. Its earlier history is reviewed in the previous work^39^. The analysis examines the magnitude and nature of the constraints, both internal and environmental. A typical application of surprisal analysis to a large cohort of patients (TCGA data) has been previously done^40^. It has been applied to time course data, for example^33^ when the magnitude of the constraints varies with time.

In the context of GBM, surprisal analysis has emerged as a powerful approach to decipher the complex cellular architectures and signaling networks embedded in single-cell transcriptomic data, often constrained by limited biological priors^31^. We extended this method to apply to spatial transcriptomic data in this study. By leveraging gene expression profiles, the method allows the system’s state to be decomposed into baseline (steady-state) behavior and a series of constraint-induced modes that can be mapped with the original spatial structure.

The input matrix was deciphered with the singular value decomposition (SVD) after the natural log transformation of the raw counts (Fig. 1B). The row is the spatial spot information, and the column is the gene feature names. Mathematically, the equation of the surprisal analysis (Fig. 1C) expresses the log-deviation of gene expression from steady state as:

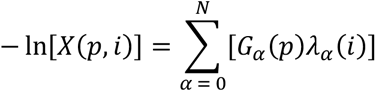

Where *X*(*p*, *i*) is the expression of gene *i* in region *p*, *λ_α_*(*i*) represents the contribution of gene *i* to the singular value *α*, and *G_α_*(*p*) quantifies the amplitude of that singular value *α* in spatial region *p*. This decomposition identifies dominant regulatory programs and spatially coherent gene modules that underlie GBM heterogeneity.

The steady state is defined by the *α* = 0 term. For each of the constraints *α* ≥ 0, the gene contribution *λ_α_*(*i*) is different, and the amplitude *G_α_*(*p*) of this constraint only differs when the spatial region *p* differs. More specifically, when *λ_α_*(*i*) and *G_α_*(*p*) have opposite signs, the product of these two contributors will add to the -ln[*X*(*p*, *i*)] in a significant way (Fig. 1D-E). The separative and cumulative terms of one specific feature (*λ_α_*(*i*)) and a certain spot shows the spatial profiles of the gene expression decomposition (Fig. 1F). The cumulative terms will ultimately approach the original feature expression. The top gene contribution *λ_α_*(*i*) with the signed value can then be divided into two groups (bottom: 5% quantile; top: 95% quantile) corresponding to the positive (red) and negative (blue) values of spatial amplitude *G_α_*(*p*).

With standard unsupervised clustering using Seurat and the surprisal analysis, we dived into these 10 patient samples with the phenotype composition and characterization of each sample based on a three-fold manual method that included i) consultation with three neuropathologists on the H&E for annotation, ii) PhenoCycler staining for cell types, and iii) manual analysis of marker gene lists defining each unsupervised cluster using Seurat. This characterization confirmed the presence of GBM subtypes (NPC-like, MES-like, AC-like), immune cells, neurons, astrocytes, oligodendrocytes, and perivascular niches. Hematoxylin and eosin staining of all ten specimens is shown in and an integrated dimensionality reduction embedding across all samples, annotated by phenotype (Fig. 1G, Supplemental Fig, 1), and demonstrate the collective GBM tumor microenvironment (Fig. 1H).

### Cross-sample surprisal analysis reveals shared spatial programs and comparison with PCA

Having established the surprisal analysis framework, we next sought to determine whether the identified constraint modes reflect shared spatial programs across the GBM cohort or are driven by individual patient variation. To address this, we performed a joint surprisal analysis across all ten samples simultaneously, enabling direct comparison of constraint modes across patients and tumor regions.

Each vector *G_α_* and *λ_α_* of different constraints *α* can be interpreted globally across the samples. The spatial amplitude of the first constraint mode, *G*_1_(*p*), was plotted across all pixel indices for the full cohort (Fig. 2A). The distribution of *G*_1_ values across samples revealed consistent partitioning of spatial spots into positive and negative amplitude groups, suggesting that the first constraint captures a recurrent spatial program present across patients and regions. To identify the biological identity of this constraint, we examined the mean gene contribution *λ*_1_(*i*) with genes organized by cell-type annotations (Fig. 2B). This analysis revealed that neuronal subtypes (cholinergic, dopaminergic, GABAergic, glutamatergic neurons), glial populations (astrocytes, oligodendrocytes), and GBM subtypes (MES-like, NPC-like, AC-like) contributed differentially to the first constraint mode, indicating that this mode captures cell-type-associated spatial variation rather than technical noise.

**Figure 2.**
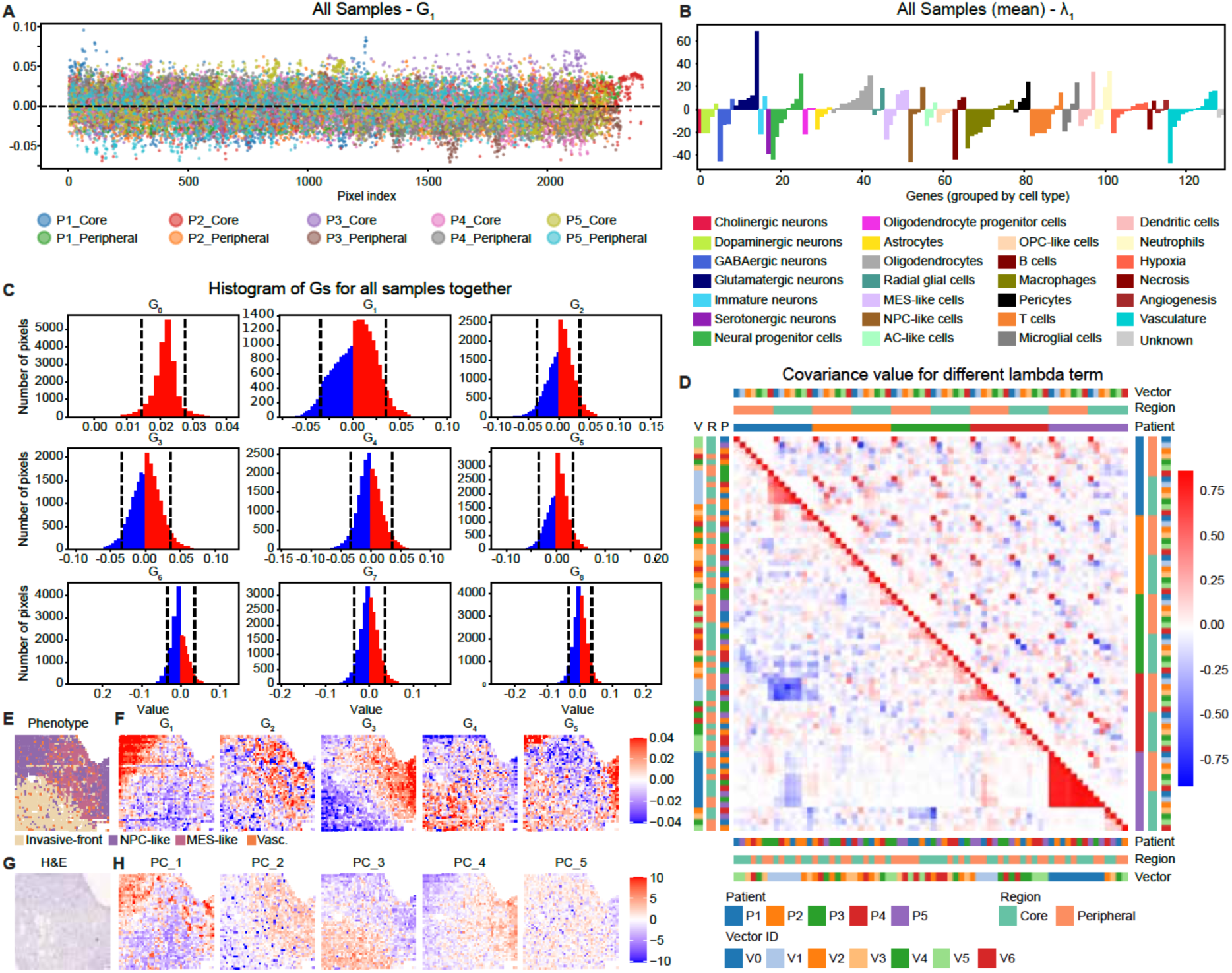
Cross-sample surprisal analysis identifies shared spatial constraint programs and comparison with PCA. (A) Spatial amplitude G1(p) of the first constraint mode plotted across all pixel indices for all ten samples (P1–P5, Core and Peripheral), with samples distinguished by color. (B) Mean gene contribution *λ*_1_(*i*) across all samples, with genes organized by cell-type annotations encompassing neuronal subtypes (cholinergic, dopaminergic, GABAergic, glutamatergic, serotonergic, immature neurons, neural progenitor cells), glial populations (astrocytes, oligodendrocytes, oligodendrocyte progenitor cells, radial glial cells), GBM subtypes (MES-like, NPC-like, AC-like, OPC-like), immune cells (macrophages, microglial cells, T cells, B cells, dendritic cells, neutrophils, pericytes), and microenvironmental signatures (hypoxia, necrosis, angiogenesis, vasculature). (C) Histograms of spatial amplitude values *G*_0_ through *G*_8_, pooled across all ten samples. The steady-state term *G*_0_displays a unimodal positive distribution, whereas higher-order constraint modes (*G*_1_–*G*_8_) exhibit approximately symmetric distributions centered near zero; 5th and 95th percentile thresholds are indicated as dashed line for each mode. (D) Covariance analysis of *λ* vectors across constraint modes, hierarchically clustered by constraint vector identity, tumor region (Core versus Peripheral), and patient origin. The heatmap reveals the degree of shared versus patient-specific or region-specific spatial programs. (E) Singular value magnitudes across constraint indices, illustrating the relative contribution and convergence behavior of each mode. (F) Spatial maps of constraint amplitudes *G*_1_ through *G*_5_ projected onto all ten tissue sections, visualizing the dominant spatial programs across the cohort. (G) Spatial maps of the first five principal components (PC_1–PC_5) for all ten samples, providing a direct comparison between PCA-derived and surprisal-derived spatial patterns. See also Supplementary Figure S2 for extended spatial constraint maps (*G*_0_–*G*_8_) and PCA maps (PC_1–PC_10) across all samples. See Methods for PCA and surprisal analysis.

Histograms of the spatial amplitude values *G*_0_ through *G*_8_, pooled across all samples, confirmed the expected structure of the decomposition (Fig. 2C). The steady-state term *G*_0_ displayed a unimodal positive distribution, consistent with its role as the baseline expression level, while higher-order constraint modes (*G*_1_- *G*_8_) exhibited approximately symmetric distributions centered near zero, with 5th and 95th percentile thresholds delineating the spatial spots most strongly associated with each mode. The singular value magnitudes decreased with increasing constraint index (Fig. 2E), confirming the expected convergence behavior and indicating that the first few constraints capture the dominant spatial programs.

To assess the degree of shared versus patient-specific spatial programs, we performed covariance analysis of the λ vectors across constraint modes, hierarchically clustered by constraint vector identity, tumor region (Core versus Peripheral), and patient origin (Fig. 2D). This analysis revealed that the lower-order constraint vectors clustered primarily by biological content rather than by patient identity, indicating that the dominant spatial programs are shared across the cohort. Higher-order constraints showed greater patient-specific variation, consistent with the expected structure in which the leading modes capture conserved biology while later modes reflect individual heterogeneity.

Using P3_Core as a representative example, we examined the spatial organization of the constraint modes in relation to the previously defined phenotype clusters (Fig. 1D and Fig. 2E). P3_Core is characterized primarily by MES-like and NPC-like GBM subtypes, together with a broad invasive region annotated as the invasive front and a prominent vascular pattern. Spatial projection of the constraint amplitudes *G*_1_ through *G*_5_ across the ten tissue sections from P3_Core revealed spatially coherent patterns, with regions of positive and negative amplitude corresponding to biologically meaningful tissue domains (Fig. 2F; Supplemental Fig. 2). Comparison with the corresponding H&E image further supported the association between these constraint patterns and the underlying tissue architecture (Fig. 2G). Although Fig. 2E–H presents P3_Core as a representative sample, similar spatial patterns were observed in the other patients, suggesting that the leading constraints reflect shared organizational features of GBM rather than sample-specific artifacts.

To directly compare surprisal analysis with conventional dimensionality reduction, we performed PCA following the Seurat v5 working flow that perform an SVD on the same dataset but with natural log transformation, variable gene selection and scaling and projected the first five principal components onto all tissue sections (Fig. 2G; Supplemental Fig. 3). While both methods identified spatially structured patterns, several key differences emerged. First, PCA, which operates on mean-centered and variance-scaled data, inherently discards the steady-state information captured by the surprisal *G*_0_ term. Second, the surprisal-derived constraint modes showed faster convergence, consistent with predictions from data compaction theorems for log-transformed data. Third, visual comparison of the spatial maps revealed that surprisal-derived constraints produced spatial patterns that more closely aligned with histologically defined tissue domains, supporting the utility of the information-theoretic framework for structured decomposition of spatial transcriptomic data. We note that both approaches identify overlapping aspects of spatial variation, as expected given their shared reliance on SVD; the distinction lies in the preprocessing and interpretive framework, with surprisal analysis offering a principled connection to the thermodynamic concept of deviations from a biological steady state and the results indicate more refined and distinct patterns even at the higher order of deposition while the PC patterns become less distinguishable with the same color scaling cap (95 percentile).

### Surprisal analysis uncovers hidden biological processes as well as conserved processes across samples

To better understand the biological processes underlying the constraint patterns identified through surprisal analysis, particularly those emerging in constraints that diverge from PCA-derived spatial patterns, we performed Gene Ontology (GO) analysis on a subset of pixels defined by the surprisal analysis. This subset captured the top 5% most positive and most negative values, represented visually by the darkest red and darkest blue respectively.

Using P1_Core as our initial sample, we selected these pixels for the GO analysis to gain insight into the biological significance of this pattern (Fig. 3A). Importantly, it is evident that the spatial patterns identified from surprisal differ from those of PCA on the same sample, with the surprisal continually revealing new, distinct, patterns with each constraint while the PCA often failed to resolve GO pathways (Fig. 3B, Supplemental Fig. 4). GO analysis results based the most negative pixels (blue) in G_3_ revealed a coherent spatial pattern that was enriched for immune-related biological processes, including immune response–activating signaling, regulation of the innate immune response, and lymphocyte differentiation (Fig. 3C). This enrichment was notable because phenotype clustering did not suggest a prominent immune compartment in this region in this spatial region but rather was characterized mainly by oligodendrocytes (Fig. 3D-E). To better spatially contextualize these signals, we spatially mapped the module score for the top-ranked immune pathway, immune response-activating signaling, which recapitulated the localized enrichment identified by surprisal analysis. We then validated this pattern using PhenoCycler multiplexed imaging (Fig. 3F). As expected, OLIG2 was broadly positive across the tissue, while CD45 and CD68 signal was concentrated in the top-right and central regions and recapitulates the general enrichment identified by surprisal. The ability to identify these localized immune-associated programs within regions not classified as immune-enriched highlights the sensitivity of surprisal analysis in detecting subtle microenvironmental states that may contribute to tumor progression and therapeutic response.

**Figure 3.**
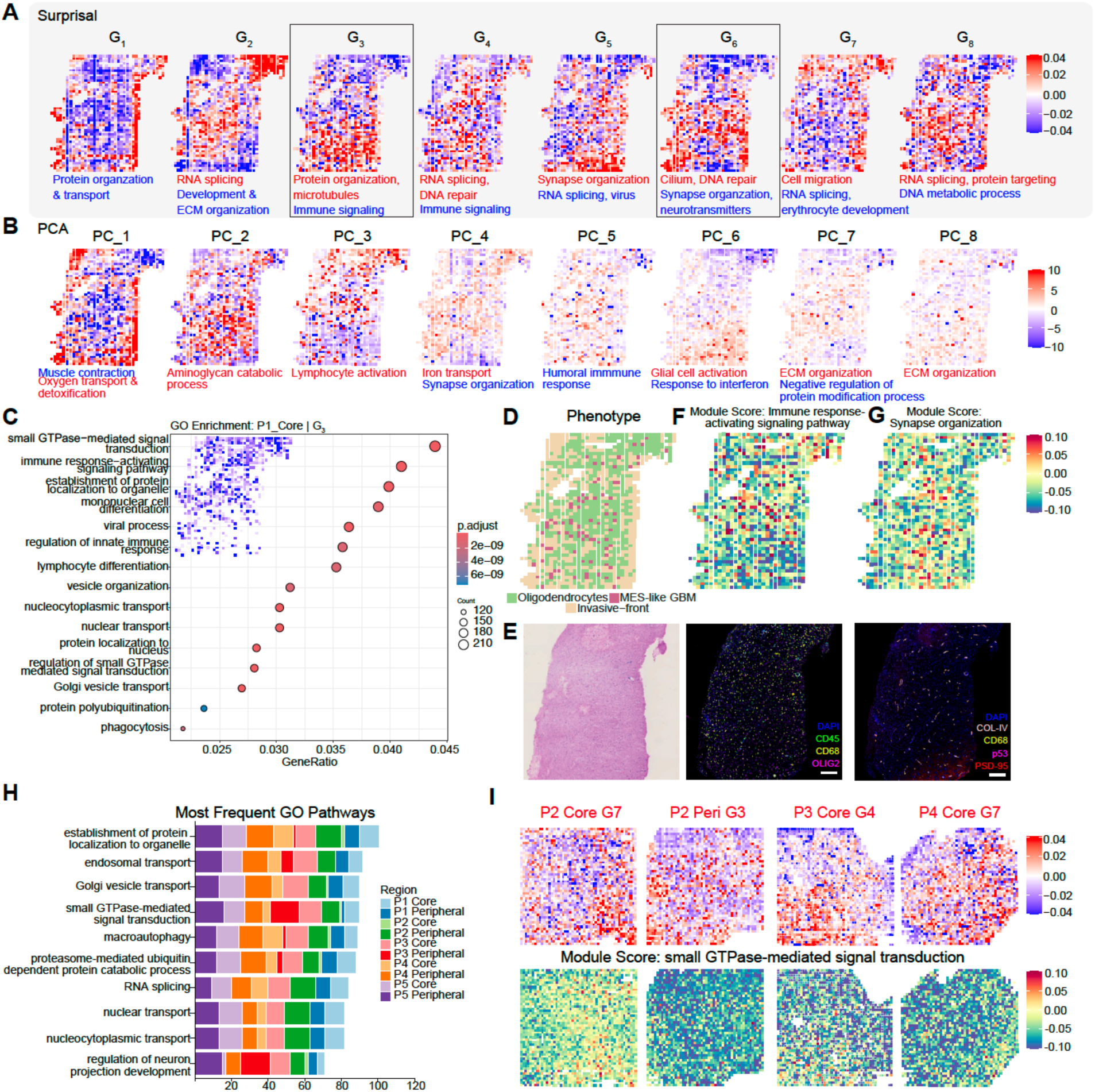
Surprisal analysis for spatial GBM uncovers both sample-specific and universal patterns. (A) Surprisal analysis of sample P1_Core *G*_1_-*G*_8_ reveals patterns not apparent in PCA (B). For each *G*, genes were identified from the upper and lower 5^th^ percentiles for GO analysis with major themes reflected. (C) GO pathway analysis for genes identified from P1_Core, *G*_3_ most negative 5^th^ percentile. (D) Phenotype analysis of sample P1_Core. (E) H&E image of sample P1_Core. (F) Expression of module score of top GO pathway identified in *G*_3_, immune response-activating signaling pathway (top) and phenocycler image of immune cells and oligodendrocytes (bottom). (G) Expression of module score of top GO pathway in *G*_6_, synapse organization (top), and phenocycler image. (H) Identification of most common GO pathways among all *G*’s of all samples. (I) Surprisal analysis and module score of various samples (positive *G*) that identified small-GTPase mediated signal transduction (negative *G*).

Additionally, surprisal analysis of the most negative pixels (blue) in G_6_ unexpectedly identified synapse organization as a top GO pathway (Supplemental Fig. 5). Notably, like immune cells, neurons were not captured as part of the phenotype clustering (Fig. 3D-E). We again spatially plotted the module score of this pathway and utilized PhenoCycler imaging. These findings confirmed the presence of PSD-95 as neuronal remodeling indicative of synapse organization (Fig. 3G). Together, focusing on the patterns identified in specific constraints enabled us to identify pathways reflecting microenvironmental cues that could promote tumor remodeling that were not readily captured otherwise.

Beyond sample-specific findings, surprisal analysis also revealed biological processes conserved across multiple samples and constraint levels (Fig. 3G). Notably, small GTPase-mediated signal transduction was recurrently identified as a top pathway across several samples, including P2_Core *G*_7_ Top, P2 Peri *G*_3_ Top, P3_Core *G*_4_ Top, and P4_Core *G*_7_ Top, with module scores confirming consistent spatial enrichment in each case (Fig. 3I), demonstrating the potential role of surprisal analysis in identifying aggressive or therapy resistant tumor regions.

Taken together, these results demonstrate that surprisal analysis can uncover both spatially unique transcriptional programs as well as conserved biological processes across heterogeneous tumor samples, providing complementary insight beyond standard phenotype clustering approaches.

### Surprisal analysis and interactypes reveal neuronal remodeling and neuron-tumor interactions at the invasive-front

Regulation of neuron projection appeared to be present as significant signaling mechanisms across several constraints in several samples (Fig. 3H). Recent studies have also pointed to the role of neurons in promoting invasion and remodeling, and there suggests an increasingly important contribution of the neuronal microenvironment on GBM progression, such as the formation of malignant neuron-to-glioma networks (Fig. 4A) and synaptic and electrical integration in neuronal circuits that promote glioma progression^23, 41^. Surprisal analysis captured distinct regions of neuron in close contact with tumor and inflammation (Fig. 4B, Supplemental Fig. 6C) that align with the PhenoCycler and phenotype clustering annotations (Fig. 4C-D, Supplemental Fig. 6C, E). GO analysis of the top 5% in the surprisal analysis of the fifth constraint for sample P1_Peripheral demonstrates the significance of synaptic remodeling and organization, and in the top 5% of the second constraint for sample P2_Core.

**Figure 4.**
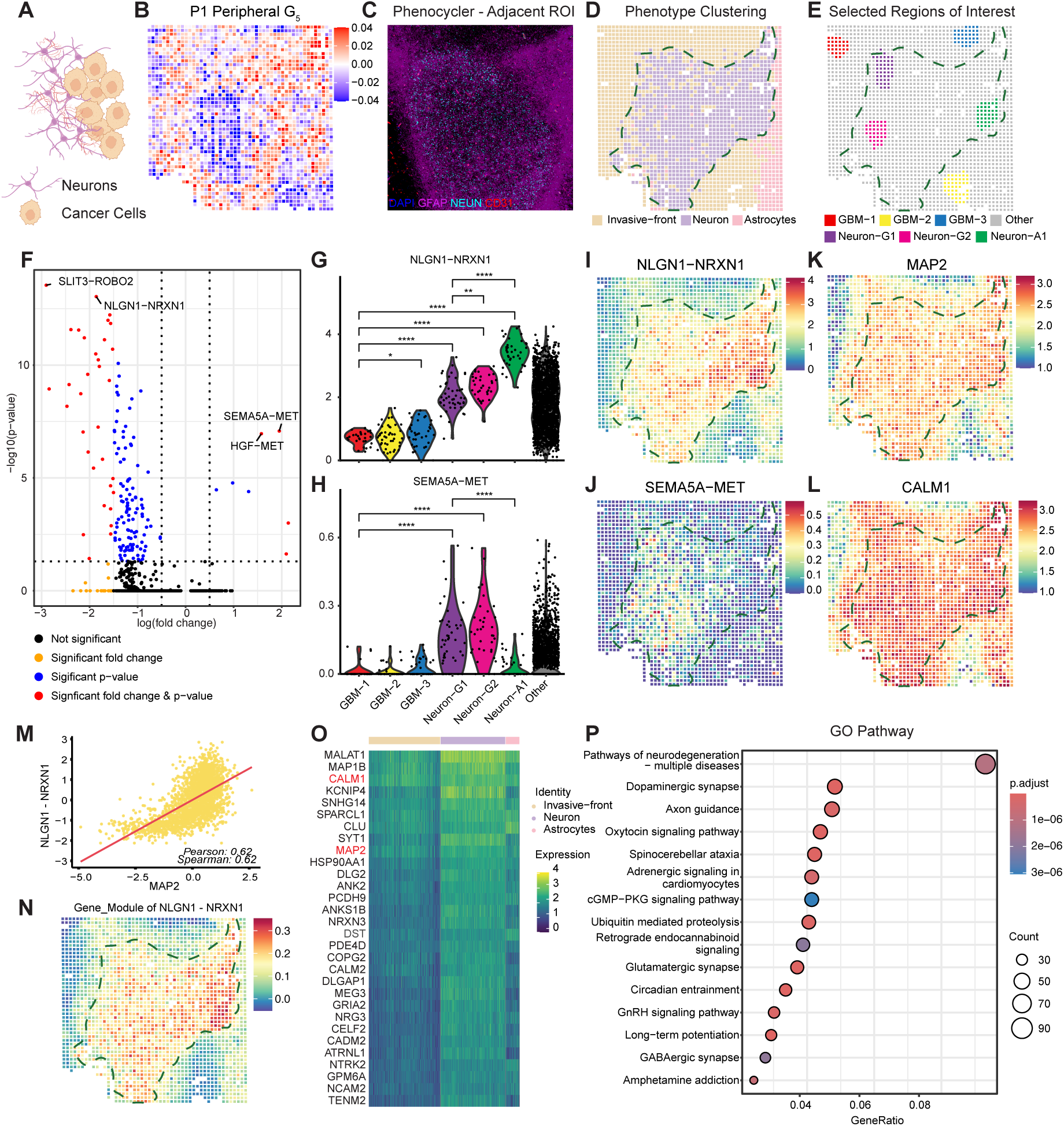
Neuronal remodeling and signaling mechanisms identified by NICHES. (A) Schematic of neuron-tumor interactions, that have become increasingly appreciated for their role in GBM progression. (B) Surprisal analysis of sample P1_Peripheral. (C) CODEX staining validates presence of neurons in the central region (NEUN+), along with immune cells and reactive astrocytes in the periphery (GFAP+). (D) Spatial phenotype clustering demonstrates adjacent invasive-front and neuronal regions. (E) Manual selection of GBM regions, neurons spatially adjacent to invasive-front (Neuron-G1, Neuron-G2), and neurons spatially adjacent to astrocytes (Neuron-A1). (F) Differentially expressed ligand-receptor interactions between neurons adjacent to GBM cells, compared to neurons adjacent to astrocytes. Significant ligand-receptor interactions have an absolute value average log2 fold change greater > 1.5, and adjusted p value < than 0.05. (G) NLGN1-NRXN1 was significantly enriched in neurons closest to astrocytes, compared to neurons closest to GBM cells. (H) SEMA5A-MET was enriched in neurons closest to GBM cells, compared to GBM cells alone or neurons close to astrocytes. (I) Spatial representation of enrichment in G. (J) Spatial representation of enrichment in H. (K-M) Pearson correlation for genes correlated with NLGN1-NRXN1 revealed MAP2 and CALM1 are positively correlated with this signaling mechanism. (N-O) Gene module and heatmap of the top 30 genes correlated with NLGN1-NRXN1 expression show specificity of these genes in neurons. (P) GO analysis of top genes correlated with NLGN1-NRXN1 signaling mechanism.

Given the dynamic nature of these pathways, we sought to understand the dominant ligand-receptor interactions that characterize these regions. We utilized a bioinformatics tool, NICHES, with the ligand-receptor database FANTOM5 to investigate the neighborhood-to-cell dynamics in the different phenotypical sub-regions. NICHES transforms conventional L-R analysis by elevating every potential interaction to single-edge resolution and embedding it directly within the spatial architecture of the tissue^42^. Starting from a standard Seurat object, it constructs sparse graphs that link each sender–receiver cell pair, optionally constrained by k-nearest neighbors or a physical radius so that only biologically plausible edges from the FANTOM5 database are kept (nine spots in this experiment). All cell-level metadata carry through unchanged, so annotations previously defined regions can be overlaid on the interaction map.

To understand the interactypes of this interface, we used Shiny to manually select regions of clusters identified as Neuron that are spatially in direct contact with clusters identified as Invasive-front, to show the potential interacting manner of neurons and cancer cells (Fig. 4E, Supplemental Fig. 6F). Differentially expressed Ligand-Receptor (LR) mechanisms between interactypes among these regions revealed NLGN1–NRXN1 as one of the top mechanisms that were highly expressed in the neuron region, a prototypical synaptic adhesion and neurotransmission LR pair (Fig. 4F-G, Supplemental Fig. 6G-H). In P2_Core, EFNA5-EPHA6 was identified as upregulated in neurons away from the bulk tumor (Supplemental Fig. 6L-M). Additionally, we found SEMA5A-MET (Fig. 4H, J, Supplemental Fig. 6J) as enriched in the neuronal regions closest to the invasive front compared to the neurons located adjacent to astrocytes. SEMA5A has been shown to interact with MET to mediate activity of HGF and suppress cell migration and invasion in GBM^43,44^.

We then calculated the gene module scores highly correlated to NLGN1–NRXN1 using the Pearson correlation and Spearman correlation methods. MAP2 (Fig. 4K) and CALM1 (Fig. 4L) were the genes top related. Neuroligins (NLGNs), such as NLGN1, reside on the postsynaptic membrane, while neurexins (NRXNs), such as NRXN1, are found pre-synaptically^45^. This LR pair is critical for synapse formation and maintenance. CALM1 (Calmodulin 1) encodes a calcium-binding messenger protein pivotal in calcium signaling, synaptic plasticity, and neurite outgrowth^46^. MAP2 (Microtubule-Associated Protein 2), a classic marker of neuronal differentiation, stabilizes microtubules in dendrites and is essential for neuronal architecture and function^47^. CALM1 and MAP2 are neuronal genes whose expression in GBM tissue provides insight into the preservation or disruption of neuronal programs in the tumor microenvironment.

We also clustered the top 30 genes correlated with NLGN1–NRXN1 expression into a heatmap (Fig. 4O) with the cell type groups showing the neuron-related gene patterns. A Gene Ontology (GO) pathway dot plot (Fig. 4P) then reflected dopaminergic synapses and GABAergic synapses that were previously investigated to play a role in GBM disease progression. Correlative analyses of CALM1 and MAP2 expression with NLGN1–NRXN1 LR activity across spatial spots revealed that the neuronal gene expression is spatially aligned with neuron–tumor interaction hotspots. This positive correlation may indicate that regions with preserved or hijacked neuronal signaling are functionally linked to ligand-receptor-mediated communication, potentially inhibiting tumor invasion, plasticity, or niche formation. Thus, examining these genes together helps illuminate a possible axis of neuron–tumor crosstalk in the GBM microenvironment.

## Discussion

Our findings uncover well-defined, spatially coherent biological programs that are not readily captured by conventional single-cell or spatial transcriptomic analyses. The identified constraint modes correspond to key processes such immune activation, and neuron–tumor synaptic interactions in human glioblastoma (GBM), demonstrating how surprisal analysis can offer additional perspectives on how microenvironmental cues influence tumor organization and behavior not captured in PCA alone. By decomposing spatial omics data into orthogonal regulatory modes, this approach reveals deviations from steady-state expression that reflect coordinated biological responses, providing a window into the hidden regulatory structure of complex tissues. To our knowledge, this represents the first integration of information-theoretic principles with spatial multi-omics analysis, underscoring the potential of such frameworks to expose latent regulatory programs that shape tumor biology beyond the resolution of traditional analytical methods.

The biological insights from this framework align with and extend recent studies on the GBM microenvironment. We demonstrate that applying surprisal analysis to spatial GBM data, we can uncover pathways common among all samples, such as small GTPase mediated signal transduction, as well as regulation of neuron projection development. At neuron–tumor interfaces, we detected ligand–receptor pairs such as NLGN1–NRXN1 and SEMA5A–MET, along with neuronal gene modules including MAP2 and CALM1, that may suggest synaptic remodeling and hijacking of neuronal communication. These findings resonate with emerging evidence that gliomas can integrate into neural circuits and exploit activity-dependent signaling to support growth and invasion. By capturing these processes within intact human tumor tissue, our study adds spatial resolution and molecular depth to the understanding of how neuronal niches influence GBM progression. Extending these analyses across other GBM niches or interfaces (Figure 1E) could enable the depiction of a complete spatial landscape of human GBM, even though individual tumors display only a subset of these niches and their underlying disease mechanisms.

There are limitations in this study that frame the interpretation of our data and results. The analysis was based on ten tissue samples from five patients, which necessarily constrains the ability to examine the association with clinical outcomes or therapeutic response although the collective analysis from these distinct samples can provide a comprehensive understanding of GBM architecture, niches, and cell-cell interactions. Each patient’s tumor presents a unique landscape, and rather than direct comparisons between patients, our approach integrates complementary spatial snapshots to assemble a more complete picture of GBM biology across niches. With this framework in place, future studies with larger and more diverse cohorts can help further validate these findings, determine their prevalence, and explore the association with patient-to-patient variability. Furthermore, while interactype analysis predicted niche-specific ligand–receptor signaling, functional studies in experimental models will help establish causal mechanisms. Looking ahead, expanding this approach to include additional modalities, such as spatial epigenomics or metabolomics, could provide a richer, multi-layered view of the tumor ecosystem. Clinically, our findings suggest that targeting cell-cell interactions within vascular and neuronal niches such as integrin-mediated adhesion or glioma-neuron synaptic coupling may represent promising therapeutic avenues. Ultimately, this study illustrates how integrating surprisal analysis with spatial omics can not only advance the mechanistic understanding of GBM biology but also help unveil potential strategies for therapeutic intervention.

## Methods

### Sample collection and preparation

Glioblastoma tumor samples for spatial transcriptomics were obtained with approval from the Institutional Review Board at Yale-New Haven Hospital from February 2019 to January 2021. All samples were collected and used for research with appropriate informed consent and with approval the Yale University Institutional Review Board (study ID 9406007680). All patients provided a signed informed consent. Two surgical specimens were collected for each surgery, and labeled by T1-Contrast Enhancing, or T2 Flair as annotated by the neurosurgeon during resection. Surgical specimens were collected immediately upon resection, embedded in OCT (Scigen OCT Compound, #4586), and snap frozen with liquid nitrogen and isopentane that was pre-chilled in liquid nitrogen. Samples were stored in –80°C until sectioning.

RNA integrity scores (RIN) were evaluated for each sample and samples with RIN scores greater than 6 were selected for DBiT. Samples were sectioned with a **Leica cryostat** at 10 um thickness in −20°C and placed in the direct center of slides for proteomic applications coated with poly-L-lysine (Epredia, #63478-AS). Forty consecutive slides were collected per sample and excess slides were stored in –80°C.

### H&E staining and imaging

H&E staining on slides adjacent to the sequenced slide was performed by Yale Histology Services. Slides were scanned using a whole slide image (WSI) scanner and digitally annotated by a neuropathologist at Yale.

### Microfluidic chips fabrication

The detailed fabrication process was described in our previous publication^48^. Standard photolithography was used to fabricate the microfluidic design of 20 µm or 50 µm- width channels on silicon wafers. The chrome photomasks were purchased from Front Range Photomasks following the design. The manufacturer’s guidelines were followed to spin-coat the SU-8-negative photoresist (SU-2010, SU-2025m, Microchem) on silicon wafers, and the featured heights were 20 µm or 50 µm, respectively. The polydimethylsiloxane (PDMS) microfluidic chips were fabricated using soft lithography. The base and curing agents were mixed thoroughly with a 10:1 ratio following the manufacturer’s guidelines (DOW SYLGARD™ 184) and poured onto the silicon wafers. The PDMS mixture was cured at 65 °C for 30 minutes after a 30-minute degassing progress. The solidified PDMS slab was extracted, and the inlets and outlets were punched for further use.

### DNA barcodes annealing preparation

For preparing annealed DNA barcode A (A1-A50, 25 µM), we mixed 10 µl each DNA barcode A (100 µM), 10 µl of ligation linker 1 (100 µM) and 20 µl of 2× annealing buffer (20 mM Tris, PH 7.5-8.0, 100 mM NaCl, 2 mM EDTA) following the annealing PCR progress. This procedure was similarly applied to annealing DNA barcode B (B1-B50, 25 µM) and ligation linker 2 (100 µM).

### Fixation and permeabilization and *in situ* polyadenylation process for Patho-DBiT

The tissues were brought to room temperature from −80 °C freezer. After 15 minutes, they were fixed with 1% freshly prepared PFA for 20 min at room temperature. The tissues were then permeabilized for 20 min at room temperature with 1% Triton X-100 (T8787, Sigma-Aldrich) in PBS. To stop permeabilization, 0.5× PBS-RI (1× PBS diluted in nuclease-free water, 0.05U/µL RNase Inhibitor) was used to clean up the tissue sections. The glass slide was air-dried and equipped with a PDMS reservoir covering the region of interest. We added 100 µL of Poly(A) wash buffer prepared by mixing 20µL 5× Poly(A) Polymerase Reaction Buffer (B0276SVIAL, New England Biolabs),), 2µL RNase Inhibitor (40,000U/mL, Y9240L, Enzymatics) and 78µl nuclease-free water into the reservoir. The tissue section was equilibrated at room temperature for 5 minutes. Poly(A) Mix made of 1× Poly(A) Reaction Buffer, 600U/µL Poly(A) Polymerase (M0276SVIAL, New England Biolabs), 100mM ATP (M0276SVIAL, New England Biolabs), SuperaseIn RNase Inhibitor (AM2694, Ambion) and Enzymatics RNase Inhibitor was added to the reaction chamber and the reservoir was sealed with parafilm. The device was Incubated in a wet box at 37°C for 25 minutes. After incubation, 0.5× PBS-RI was used to clean up the tissue sections three times.

### Bulk tissue reverse transcription

We first prepared a reverse transcription mix with 12.5 µL of 5× RT Buffer, 8.2 µL of RNase-free water, 0.4 µL Enzymatics RNase Inhibitor, 0.8 µL µL SuperaseIn RNase Inhibitor, 3.1 µL of 10 mM dNTPs each (ThermoFisher), 6.2 µL of Maxima H Minus Reverse Transcriptase (ThermoFisher), 25 µL 0.5× PBS-RI and 10 µL RT primer (/5Phos/CATCGGCGTACGACTTTTTTTTTTTTTTTTVN). The 60µL RT Mix was loaded on the sample into the PDMS reservoir and sealed with parafilm. Then, it was incubated in a wet box at room temperature for 30 minutes and 42°C for 90 minutes. To remove excessive RT primer, after the RT reaction, the slide was first dipped in 50mL PBS and then transferred and merged into a new 50mL Falcon tube with nuclease-free PBS, shaken at room temperature for 5 mins. Then, the tissue was flow-washed with 1mL of nuclease-free water twice to remove residual salt, followed by air drying.

### In situ DNA barcode ligation

In the barcode A in situ ligation process, the region of interest was covered with a PDMS slab and imaged using a 10× objective with EVOS FL Auto microscope. The PDMS device was clamped on the slide with an acrylic clamp. A ligation reaction solution was prepared with 2 µL of ligation mix (26 µL 10× T4 ligase buffer, 61.3 µL nuclease-free water, 5 µL 5% Triton-X100, 2µL Enzymatics RNase Inhibitor, 0.7 µL SuperaseIn RNase Inhibitor), 2 µL of 1× NEBuffer 3.1, and 1 µL of each annealed DNA barcode A, and loaded into 50 channels. The chip was incubated in a wet box at 37°C for 30 minutes and washed with 1× NEBuffer 3.1 thrice. Then the clamp and PDMS were removed. Finally, the slide was rinsed in water and air-dried.

In the barcode B in situ ligation process, a second PDMS slab with perpendicular channels was attached to the air-dried slide, and a bright-field image was captured using a 10× objective with EVOS FL Auto microscope. The slide was then clamped with PDMS. The preparation of the ligation mix and loading process of DNA barcode B are the same as the DNA barcode A preparation. The chip was incubated in a wet box at 37°C for 30 minutes and washed with 0.5× PBS-RI thrice. The PDMS chip was peeled off, and the slide was dipped into a 50 mL tube filled with 0.5× PBS-RI first, then transferred into a new 50 mL tube with 0.5× PBS-RI and shaken for 5 mins. Then, to remove residual salt, wash with 1 mL of nuclease-free water twice. The slide was air-dried for the next steps.

### Tissue lysis collection

The region of interest was covered with a clean PDMS reservoir, and the tissue and the reservoir were clamped together. The 70 µL lysis mix consisting of 35 µL 1× PBS, 35 µL 2× lysis buffer and 7 µL Proteinase K solution (20 mg/mL, EO0491, Thermo Scientific) was loaded into the PDMS reservoir and sealed with parafilm. Then, incubated in a wet box at 55°C for 2 hours to reverse formaldehyde crosslinks. The final tissue lysate was collected into a 200 µL PCR tube.

### Purification of cDNA, template switch and library establishment with rRNA removal

The lysate was purified using the DNA Clean & Concentrator-5 kit (D4014, Zymo) according to the user manual and eluted in 150 µL of nuclease-free water. We then washed the 40 µL/sample of Dynabeads Streptavidin trice with 1mL of 1× B&W buffer with 0.05% Tween-20 and resuspended the beads with 150 µL 2× B&W buffer. The beads were mixed with cDNA elution and agitated at room temperature for 60 minutes. A magnet was used to separate beads and supernatant in the lysate and the beads were washed twice with 1× B&W buffer with 0.05% Tween-20 and twice with 10 mM Tris-HCl and 0.1% Tween-20. Streptavidin beads with bound cDNA molecules were then resuspended in TSO Mix (40 µL 5× Maxima RT buffer, 40 µL 20% Ficoll PM-400, 20 µL 10mM dNTPs, 10 µL Maxima H Minus Reverse Transcriptase, 5 µL Enzymatics RNase Inhibitor, 10 µL 100 mM TSO primer: 5’-AAGCAGTGGTATCAACGCAGAGTGAATrGrG+G-3’, 75 µL nuclease-free water). The beads were incubated at room temperature for 30 minutes and 42°C for 90 minutes with gentle rotation. The beads were resuspended in a PCR mix consisting of 100 µL 2× HiFi HotStart Master Mix, 84 µL nuclease-free water, 8 µL PCR Primer 1 (10 µM, 5’-CAAGCGTTGGCTTCTCGCATCT-3’), 8 µL PCR Primer 2 (10 µM, 5’-AAGCAGTGGTATCAACGCAGAG-3’) after washing beads using 10 mM Tris with 0.1% Tween-20 once and water once. The PCR process followed a thermocycler program: 95°C for 3 min and cycling at 98°C for 20 s, 65°C for 45 s and 72°C for 3 minutes for five times, then 72°C for 5 minutes. The Dynabeads were removed by a magnet with 5 mins binding after the PCR amplification and 1 µL 20× EvaGreen (#31000, Biotium) was added to a 19 µL PCR solution aliquot and the mixture was placed in qPCR machine (Bio-Rad) with the following thermocycling conditions: 95°C for 3 minutes, cycling at 98°C for 20 seconds, 63°C for 45 seconds, 72°C for 3 minutes for 20 times, followed by a single 5 minutes at 72°C after cycling and 4°C hold. The remaining samples underwent the same PCR program with the cycle numbers decided by 1/3 of the saturated signal in qPCR results.

The 0.8× SPRI beads (B23317, Beckman Coulter, Inc.) were used to purify the samples following the standard manufacturer’s guidance. Two rounds of rRNA removal using SEQuoia RiboDepletion Kit (17006487, Bio-Rad) were performed under the manufacturer’s guidance. For different tissue samples, pivot experiments need to be done, and additional rounds of depletion are recommended.

After the rRNA depletion step, another 10 cycles of PCR reaction using a 100uL PCR reaction system (50 µL 2× Kapa HiFi Master Mix, 10 ng cDNA, 4 µL 10 µM P7 primer, 4 µL 10 µM P5 primer, nuclease-water) were performed following the PCR process described before. We applied the 0.8× SPRI beads selection again to remove the excessive primers and eluted the cDNA to 20 µL nuclease-free water.

### Sequencing and quality control

The size and concentration of the sequencing library were assessed using an Agilent 4150 TapeStation and D5000 DNA ScreenTape and reagents (5067-5589, Agilent). Next-generation sequencing (NGS) was then performed on the Illumina NovaSeq 6000 sequencer, utilizing a paired-end, 150-base-pair mode, generating approximately 100 million read pairs per sample.

### Read preprocessing

Raw sequencing reads were processed using CUTadapt to trim adapter sequences and low-quality bases from both ends of the reads. Subsequent preprocessing was performed with zUMIs, which included the extraction of unique molecular identifiers (UMIs), read demultiplexing, and barcode assignment. Processed reads were aligned to the reference genome using STAR (Spliced Transcripts Alignment to a Reference) with default parameters optimized for spatial transcriptomics data.

### Post-processing, gene filtering and spatial tissue spot assignment

To reduce technical noise and mitochondrial contamination, genes encoded by the mitochondrial genome (MT genes) were identified and removed from the dataset prior to downstream analysis. Only barcoded spots corresponding to tissue-covered regions were retained for analysis. Tissue detection was performed using AtlasXomics Browser with the white-field scan to generate the image metadata.

### PCA, unsupervised clustering, and surprisal analysis

Spatial gene expression data were processed using the Seurat v5 workflow. Gene expression levels were normalized using the ‘LogNormalize’ method, which normalizes feature counts for each pixel by the total expression, applies a scaling factor, and performs log-transformation. Highly variable features were identified for downstream analysis, followed by linear dimensionality reduction using principal component analysis (PCA) with the ‘RunPCA’ function. The number of principal components retained for subsequent analyses was selected based on the variance explained by each component, as assessed using an Elbow plot.

Pixels were then embedded into a shared nearest neighbor (SNN) graph using the ‘FindNeighbors’ function based on Euclidean distances in PCA space. Unsupervised clustering was performed with the ‘FindClusters’ function using a modularity optimization-based algorithm to identify spatially distinct transcriptional domains. For visualization of transcriptional heterogeneity, nonlinear dimensionality reduction was conducted using Uniform Manifold Approximation and Projection (UMAP) through the ‘RunUMAP’ function. Cluster-specific differentially expressed genes (DEGs) were identified using the ‘FindMarkers’ function by comparing gene expression profiles between pixel clusters.

Surprisal analysis was performed using the raw gene expression count matrix. The entries, individual gene expression profiles, were recast as their logarithmic value. The resulting matrix was decomposed using singular value decomposition (SVD) to identify major transcriptional constraints underlying spatial expression patterns. Each constraint consists of a spatial map describing its distribution across pixels and corresponding gene weights representing the contribution of individual genes. The dominant constraints, typically few, were selected based on the magnitude of their singular values and used to reconstruct a spatial gene expression pattern for each constraint. The reconstructed profile is the sum of the contributions of the dominant constraints. The accuracy is evaluated by quantitatively comparing the recovered expression profiles with the original gene expression data. Surprisal analysis was implemented using custom Python scripts available in the GitHub repository.

### Ligand-receptor interaction with NICHES

To investigate ligand–receptor interactions, we applied the NICHES framework to each subsetted dataset using the log-normalized expression matrix (Spatial) as input. NICHES was run with CellToCellSpatial, CellToNeighborhood, and NeighborhoodToCell modes enabled, using the FANTOM5 ligand–receptor database of the human species, the “CellType_GLSB_manual” cell annotation, and a neighborhood size of k = 9.

To analyze cell–cell signaling patterns based on the NeighborhoodToCell_Spatial assay produced by NICHES, we performed unsupervised clustering: variable feature selection, scaling, PCA (100 components), neighborhood graph construction, and Louvain clustering (resolution = 0.5). A UMAP embedding was generated using the top 30 principal components. Cluster-specific ligand-receptor signatures were identified using *FindAllMarkers* with a minimum expression threshold (min.pct = 0.5) and log-fold change threshold (logfc.threshold = 0.25). The top 10 and top 30 marker genes per cluster were ranked by average log2 fold-change and exported as CSV files. *SpatialFeaturePlots* were generated for all top marker genes across spatial coordinates with image background suppressed, to visualize the spatial distribution of key ligand–receptor signals.

### PhenoCycler for the multi-plex proteomics

PhenoCycler-Fusion imaging was performed per the standard manufacturer’s protocol (Akoya Biosciences, https://www.akoyabio.com/wp-content/uploads/2021/01/CODEX-User-Manual.pdf). Briefly, fresh-frozen tissue sections (10 µm thickness) were mounted onto positively charged microscope slides. Slides were fixed in acetone for 10 minutes and 1.6% PFA for 10 minutes at room temperature followed by air drying and rehydration in the rehydration buffer. Slides were then incubated with a blocking solution provided by Akoya Biosciences for 30 minutes. Tissue sections were then incubated at room temperature for 3 hours with a panel of oligonucleotide-conjugated primary antibodies (Akoya Biosciences and in-house conjugated using Lightning-Link® Streptavidin Antibody Labeling Kit) targeting biomarkers of interest. Following incubation, slides were washed thoroughly and then post-fixed using 1.6% PFA and treated with ice-cold methanol before a final fixative step.

The PhenoCycler imaging process used iterative cycles of fluorescent dye-conjugated oligonucleotide reporters (Akoya Biosciences) to hybridize and visualize bound primary antibodies. Each imaging cycle included reporter dye hybridization, image acquisition, and subsequent dye removal to prepare slides for the next cycle. Images were acquired using an inverted fluorescence microscope with a 20X objective.

## Data and Code Availability

Raw and processed Patho-DBiT data for all patient samples have been deposited in GEO. Raw PhenoCycler data have been deposited in Zenodo. Scripts for data analysis and visualization were written primarily in Shell, Python, and R.

## Acknowledgments

We thank Yale West Campus cleanroom for assistance with microfluidics device fabrications, the Yale Center for Genome Analysis (YCGA) for sequencing service, the Yale Pathology Tissue Services team for FFPE tissue sectioning, and the Yale Center for Research Computing for use of the research computing infrastructure. Illustrations were created with BioRender.com.

## Author Contributions

Experimental techniques: SB, GL. Data acquisition: SB, GL, MZ, KA, JM. Data analysis: SB, GL, SG, NW, HL, LM, YL, JS, ZT, WE, DZ, MD, MT, MY, KL, MSBR, FR, RDL, JZ, RF. Data curation: SB, GL, SG, NW, PG, DM, ZEO. Project administration: JRH, MSBR, FR, RDL, JZ, RF. Writing: SB, GL, SG, NW, MSBR, FR, RDL, JZ, RF.

