## Supplemental Table 1, Supplemental Figure 1, Supplemental Figure 2, Supplemental Figure 3, Supplemental Figure 4, Supplemental Figure 5 for "Decoding Spatial Programs in Human Glioblastoma Through Surprisal Information-Theoretical Analysis"

### Supplemental Materials

| Tumor identification | Official Diagnosis | IDH status | TERTp mutation | EGFR amplificaiton | Tumor Location |
|---|---|---|---|---|---|
| Patient 1 Core | Recurrent Glioblastoma, WHO Grade IV | IDH1 (R132H) | Yes at 1295228 | No | Left Temporal Insular |
| Patient 1 Peripheral | | | | | |
| Patient 2 Core | Recurrent Glioblastoma, WHO Grade IV | IDH1 (R132H) | Yes at 1295228 | No | Right Frontal Lobe |
| Patient 2 Peripheral | | | | | |
| Patient 3 Core | Glioblastoma, WHO Grade IV | IDH1 (R132H) | Yes at 1295228 | No | Left Temporal Lobe |
| Patient 3 Peripheral | | | | | |
| Patient 4 Core | Glioblastoma, WHO Grade IV | IDH1 (R132H) | Yes at 1295250 | Yes | Left multifocal tumor |
| Patient 4 Peripheral | | | | | |
| Patient 5 Core | Recurrent Glioblastoma, WHO Grade IV | IDH1 (R132H) | Yes at 1295250 | Yes | Right Temporal Tumor |
| Patient 5 Peripheral | | | | | |

**Supplemental Table 1. Patient Metadata and Cell Type Annotation.** (A) Patient information for sequenced samples. (B) Cell type characterization for all samples.

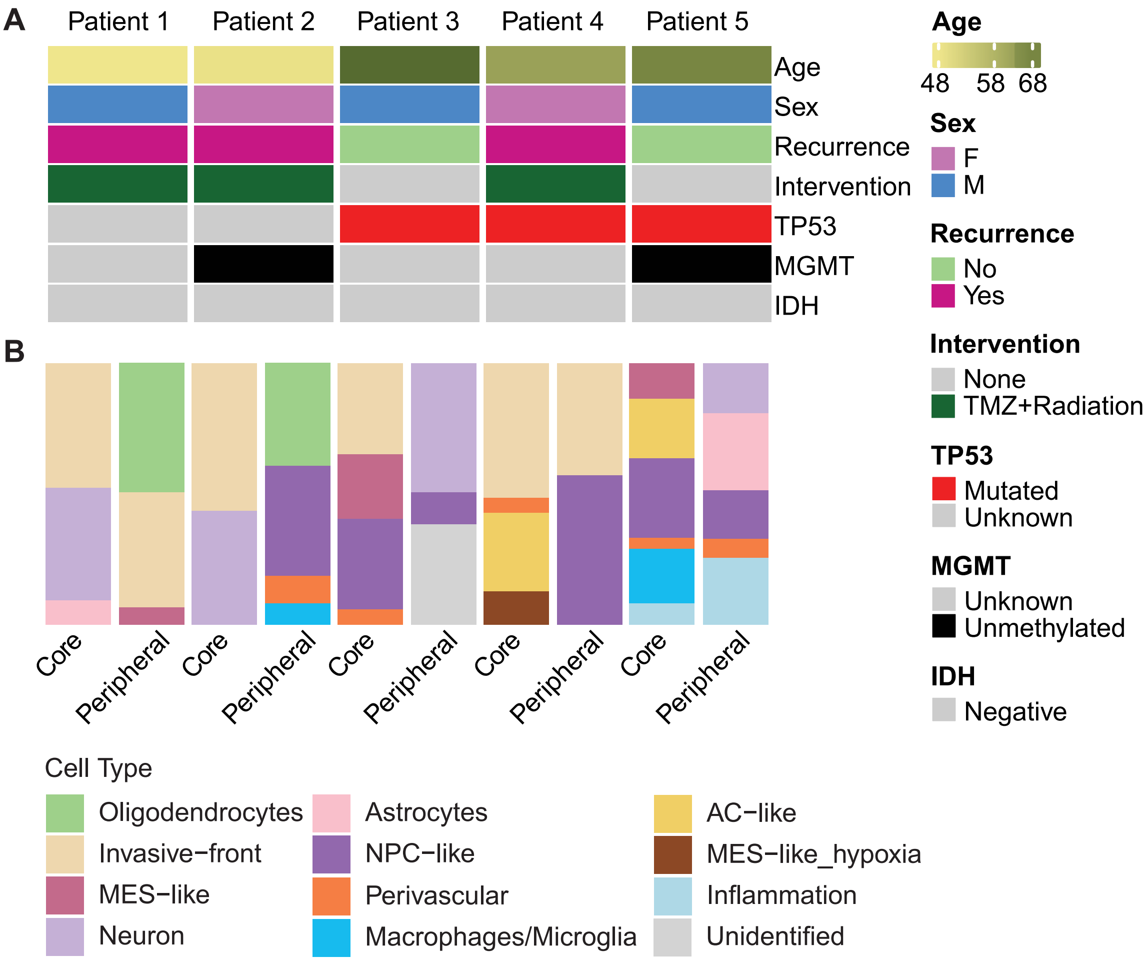


**Supplemental Figure 1. Sample information and characterization.** (A) Patient information including age, sex, recurrence status, intervention, and TP53, MGMT, IDH mutation status. (B) Phenotype clustering based on pathologist annotations, Seurat unsupervised clustering, and Phenocylcer imaging.

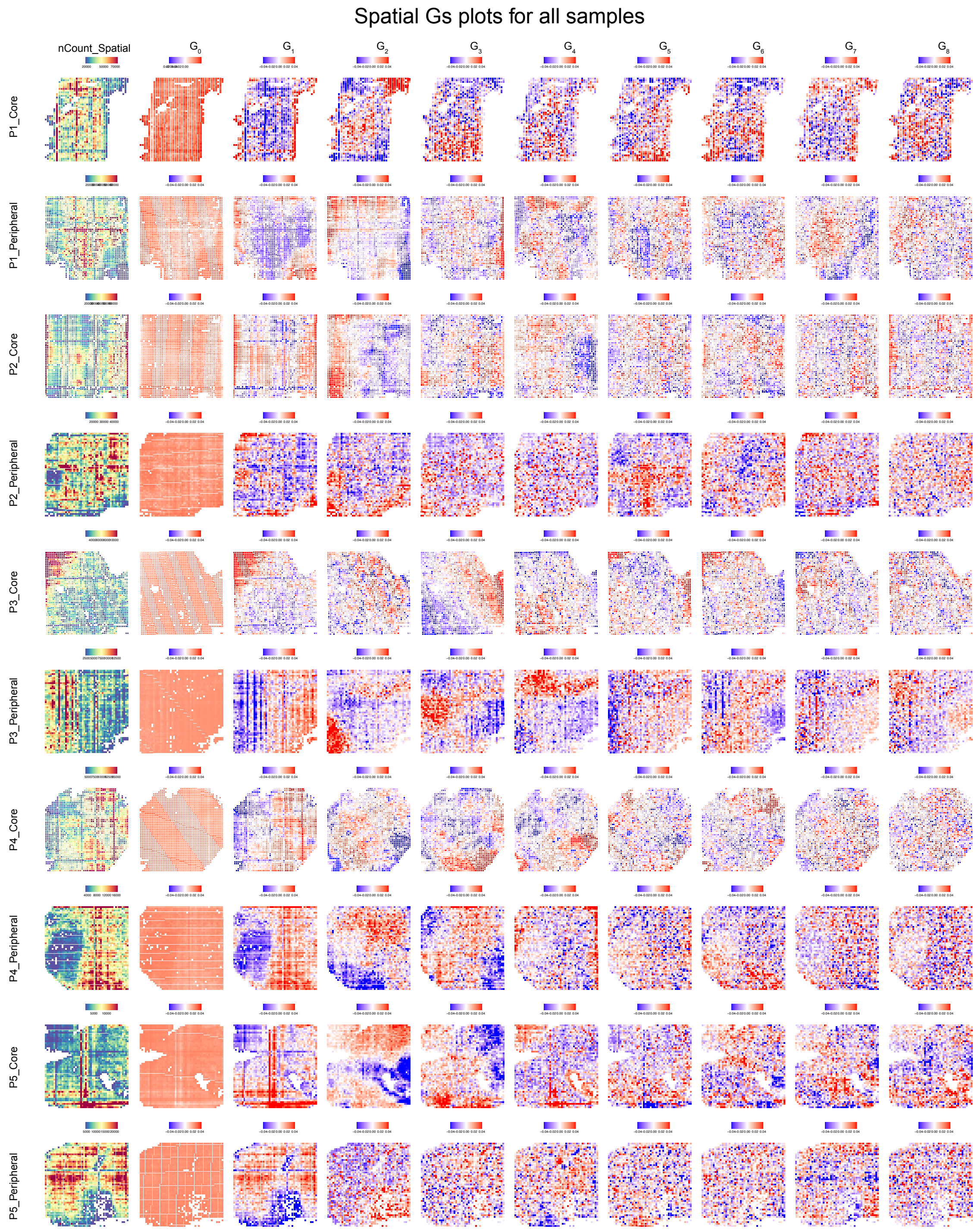


**Supplemental Figure 2. Surprisal analysis for all spatial transcriptomic samples.** The count depth (right) is shown for each sample, along with the steady state ($G_0(p)$) and subsequent constraints, $G_1(p)$ - $G_8\,(p)$.

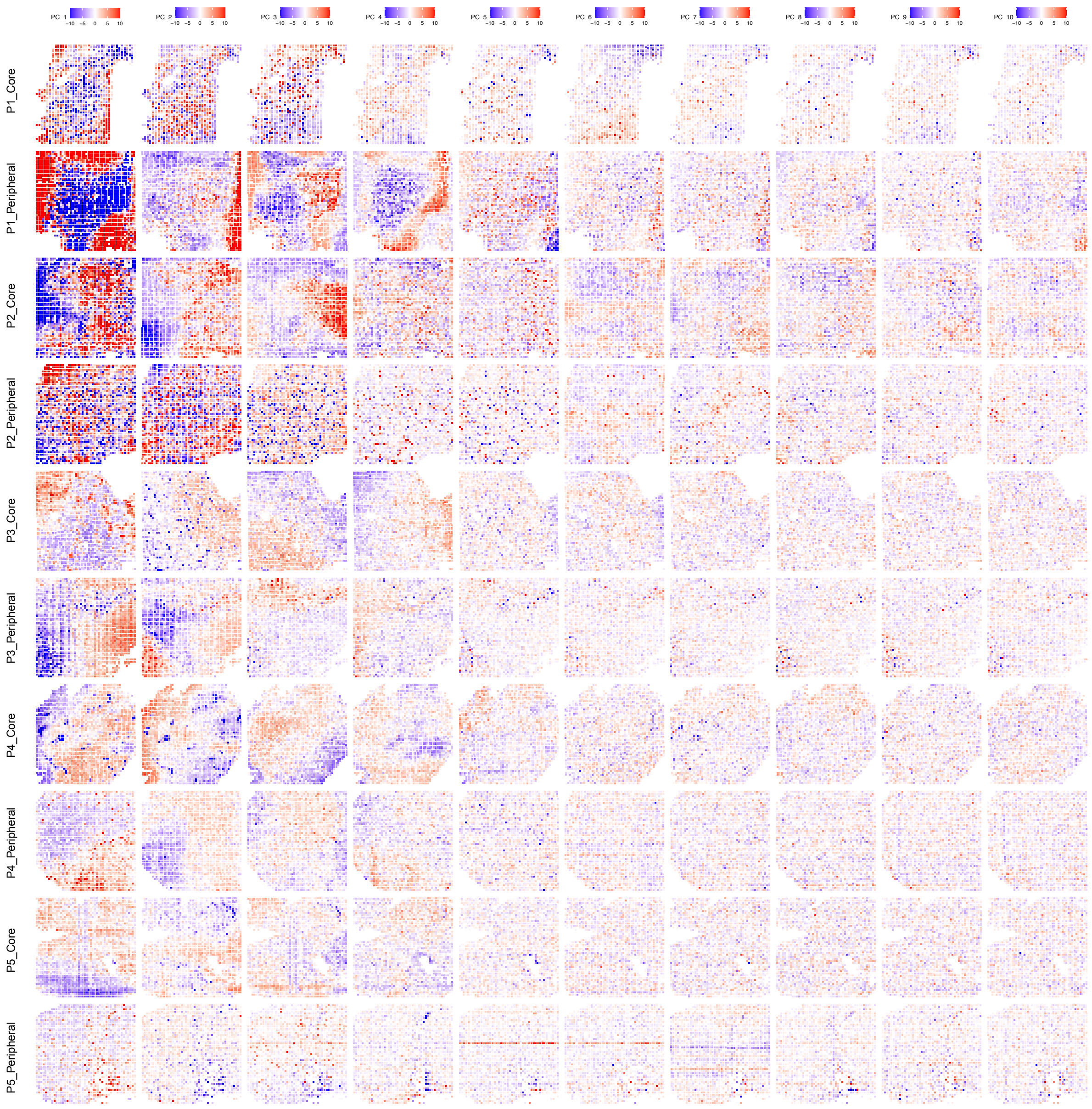


**Supplemental Figure 3. PCA for all spatial transcriptomic samples.** Eachprincipal component is shown spatially through PC_10.

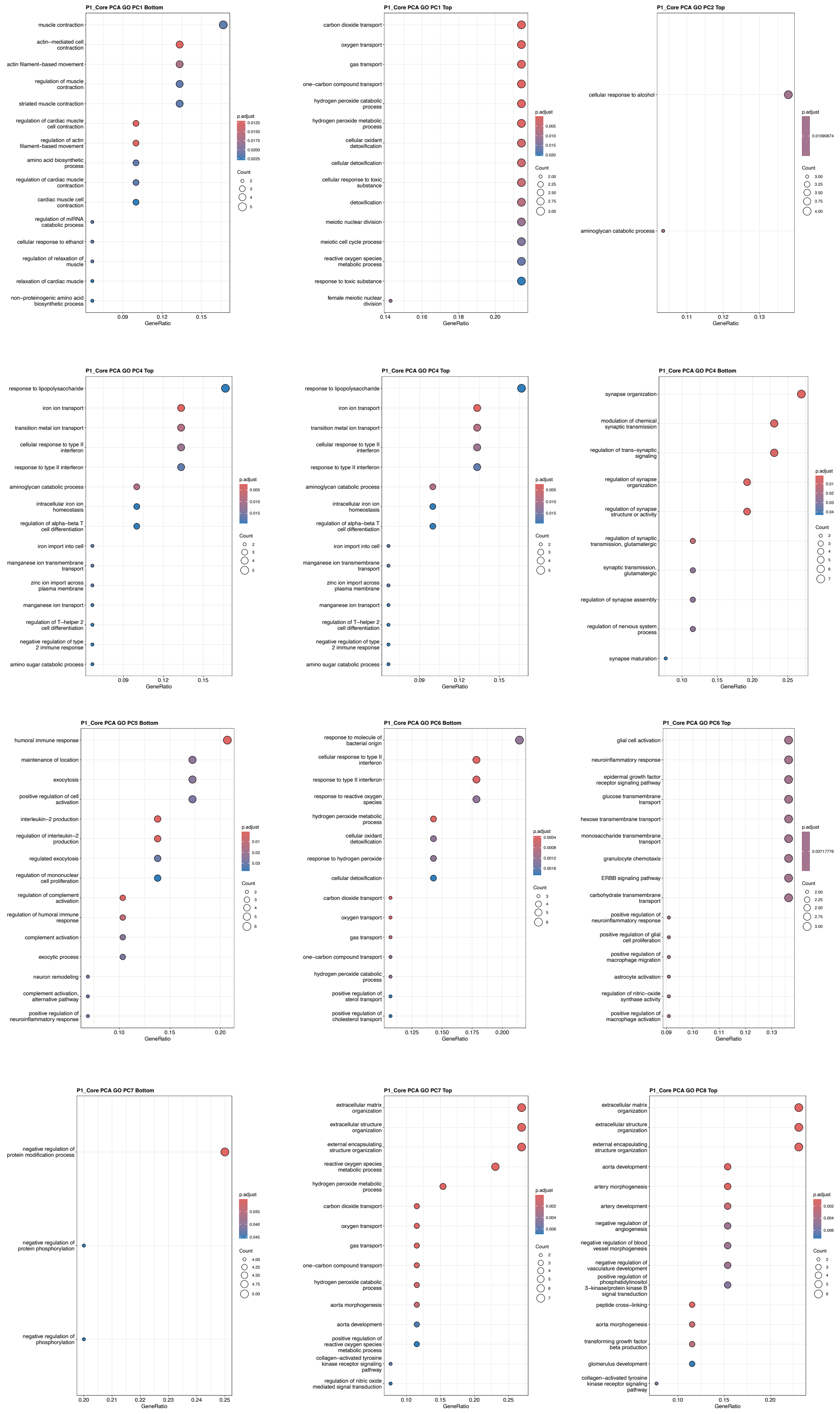


**Supplemental Figure 4. GO analysis from PCA results for P1_Core.**

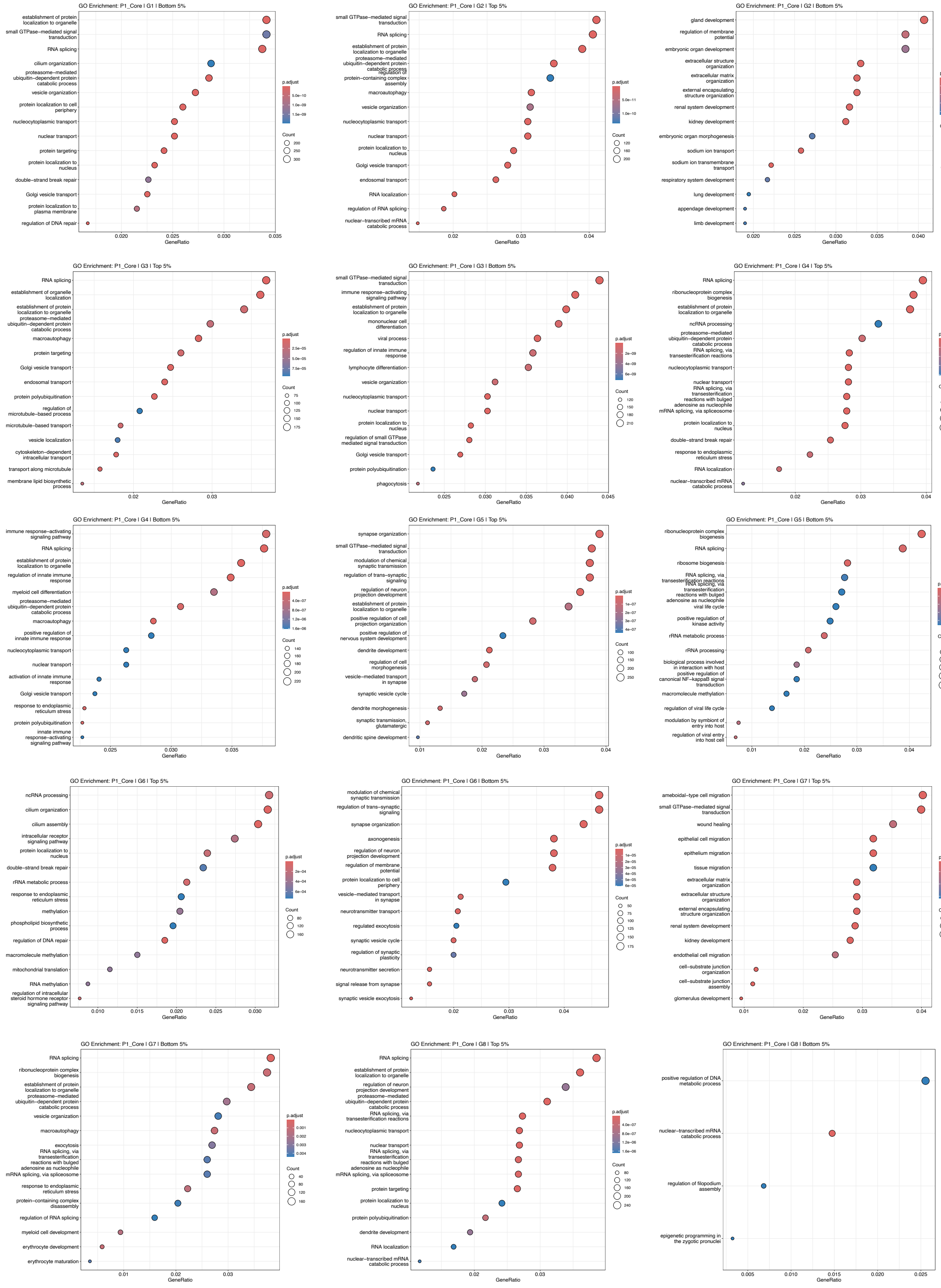


**Supplemental Figure 5. GO analysis for top and bottom 5th percentile of each constraint for P1_Core from surprisal analysis.**

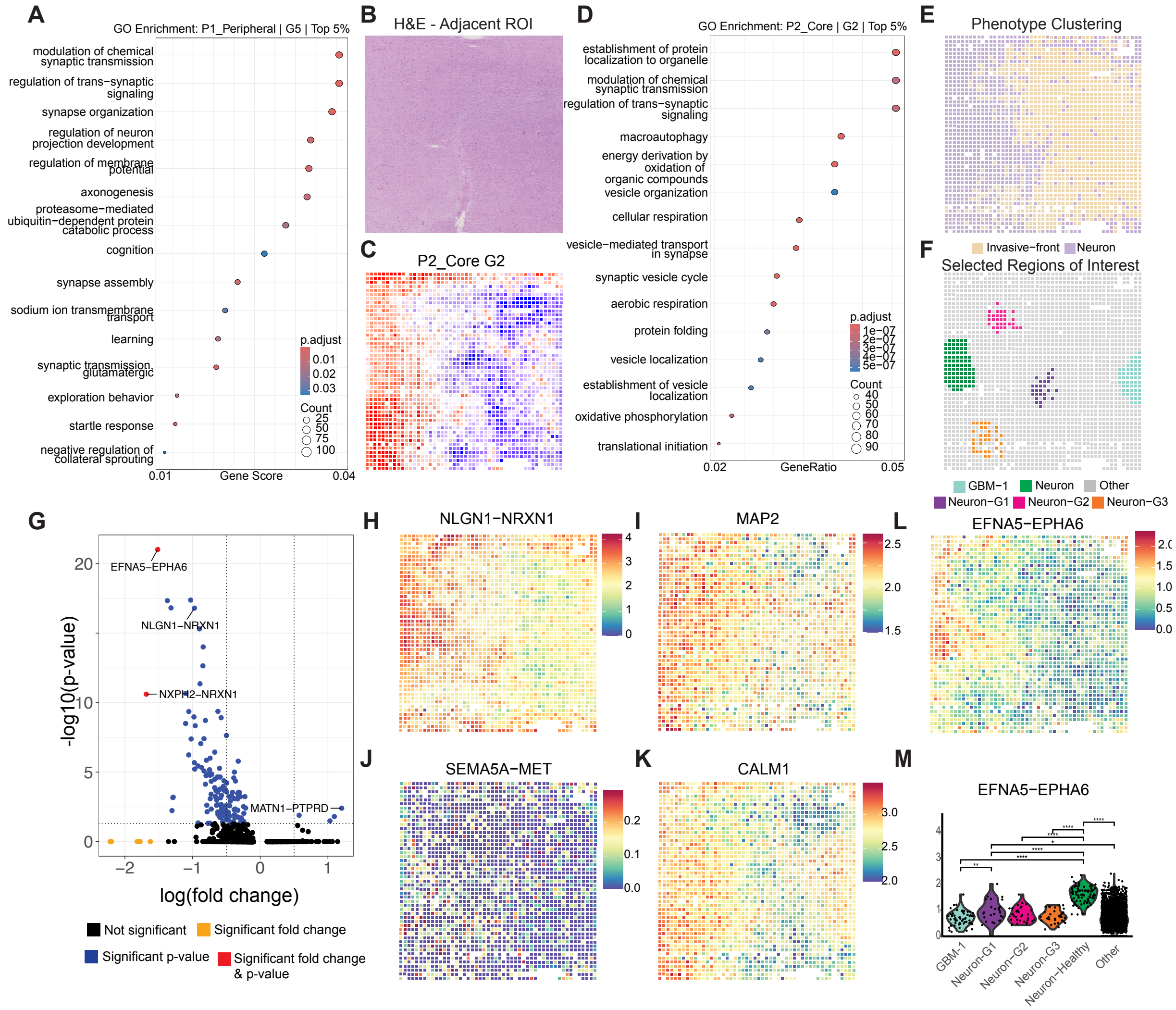


**Supplemental Figure 6. Surprisal analysis of P1_Peripheral and interactype analysis in P2_Core demonstrate neuronal remodeling.** (A) GO pathway enrichment for surprisal analysis of P1_Peripheral (red) demonstrating enrichment of pathways relevant to neuronal remodeling. (B) H&E of adjacent section of sample P2_Core. (C) Spatial pattern of G2 of surprisal analysis for P2_Core. (D) GO pathway enrichment for P2_Core top (red) demonstrating pathways relevant to synaptic signaling. (E) Spatial phenotype clusters of P2_Core demonstrate adjacent neurons and invasive-front. (F) Manual selection of GBM regions in the invasive front (GBM-1), neuron regions adjacent to invasive front (Neuron-G1/G2/G3), and neurons not spatially adjacent to invasive-front (Neuron). (G) Differentially expressed ligand-receptor interactions between neurons spatially adjacent to invasive-front, and neurons spatially near other neurons. Significant ligand-receptor interactions have an absolute value average log2 fold change greater > 1.5, and adjusted p value < than 0.05. (H) NLGN1-NRXN1 was significantly enriched in neurons closest to other neurons, compared to neurons closest to GBM cells. (I) SEMA5A-MET spatial plot showing expression in neurons furthest from neurons adjacent to tumor. (J-K) Spatial plot and quantification of EFNA5-EPHA6 that was significantly enriched in neurons furthest from the invasive-front. (L-M) MAP2 and CALM1 have a correlative spatial pattern with NLGN1-NRXN1 in P2_Core.
